# A Novel Gene *REPTOR2* Activates the Autophagic Degradation of Wing Disc in Pea Aphid

**DOI:** 10.1101/2022.09.19.508482

**Authors:** Erliang Yuan, Huijuan Guo, Weiyao Chen, Bingru Du, Yingjie Mi, Zhaorui Qi, Yiyang Yuan, Keyan Zhu-Salzman, Feng Ge, Yucheng Sun

## Abstract

Wing dimorphism is an evolutionarily adaptive trait to maximize insect fitness under various environments, by which the population could be balanced between migration and reproduction. Most studies concern the regulatory mechanisms underlying the stimulation of wing morph in aphids, but relatively little research addresses the molecular basis of wing loss. Here, we found that the wing disc of wingless-destined pea aphids degenerated 30 h post birth by autophagic rather than apoptotic degeneration, whereas winged-destined aphids developed normally. Activation of autophagy in 1^st^ instar nymphs reduced the proportion of winged aphids, and suppression of autophagy increased the proportion. The *REPTOR2* gene associated with TOR signaling pathway was identified by RNA-seq as a differentially expressed gene between the two morphs, with higher expression in the thorax of wingless-destined aphids. Further genetic analysis indicated that *REPTOR2* could be a novel gene derived from a gene duplication event exclusively in pea aphid on autosome A1 but translocated to the sex chromosome. Knockdown of *REPTOR2* reduced autophagy in the wing disc and increased the proportion of winged aphids. In agreement with REPTOR’s canonical negative regulatory role of TOR on autophagy, winged-destined aphids had higher TOR expression in the wing disc. Suppression of TOR activated autophagy of the wing disc and decreased the proportion of winged aphids, and vice versa. These results revealed that the TOR signaling pathway controlled degradation of the wing disc in pea aphids, and that REPTOR2 could modulate this autophagic degradation.

## Introduction

Flight ability is a great evolutionary success for animals that favors seeking new habitats, mates, and food resources (*Roff, 1990*). The loss of such ability has occurred in numerous insects (*Roff, 1990; Harrison, 1980; Roff and Fairbairn, 1991*). The wingless morph has evolved to optimize resource allocation from dispersal to reproduction, which facilitates the population expansion under suitable conditions (*Zera and Denno, 1997; Mole and Zera, 1993*). Alternation of the wing morph in aphid offspring is strongly influenced by maternal experience (*Braendle et al., 2006*). Pea aphids are recognized as a compelling laboratory model to study regulatory mechanisms underlying wing dimorphism, most notably through maternal population density (herein refers to as maternal density). It has been shown that ecdysteroids, miR-9b-ABCG4-insulin signaling, and two laterally transferred viral genes (*Apns-1* and *Apns-2*) are responsible for producing winged offspring in parthenogenetic females (*Vellichirammal et al., 2017; Shang et al., 2020; Parker and Brisson, 2019*). Nevertheless, the phylogenetic evidence suggests that winged morph in aphids is a default developmental pattern while the wingless morph is secondarily derived, as aphid ancestors are winged (*Braendle et al., 2006; Miyazaki, 1987*). Thus, how wingless morph is controlled under favorable conditions needs to be studied, as such information will help to elucidate the differentiation of wing dimorphism in aphids.

Tissue degradation is a common developmental process in insects executed by programmed cell death (PCD), such as autophagy and apoptosis (*Xu et al., 2019*). During metamorphosis of Drosophila, salivary gland, midgut and other larval tissues are removed by PCD (*Xu et al., 2019*). For example, suppression of apoptosis or autophagy or both could delay the degeneration of salivary glands in Drosophila (*Berry and Baehrecke, 2007*). The midgut removal during metamorphosis is severely interfered by suppression of autophagy rather than apoptosis (*Denton et al., 2012*). These findings suggest that tissue-specific degeneration required different types of PCD. Furthermore, histological examinations in pea aphids show that the 2^nd^ instar nymphs of the winged-destined morph obviously possess a pair of wing discs, whereas wingless-destined morph does not (*Ishikawa et al., 2008; Ogawa and Miura, 2013*). Since the wing bud and full wing are developed from the wing disc, it would be interesting to know whether and how PCD controls the fate of the wing disc cells, ultimately resulting in the wingless morph.

The evolutionarily conserved nutrient-sensing TOR (target of rapamycin) pathway has been implicated in the regulation of fecundity, life span as well as tissues degradation of insects in response to nutritional conditions (*Xu et al., 2019; Katewa and Kapahi, 2011*). Autophagic degradation in different insect tissues including salivary gland and midgut requires down-regulation of TOR signaling pathway (*Berry and Baehrecke, 2007; Denton et al., 2012*). In Drosophila, suppression of TOR by growth arrest activates autophagy via the Apg1 protein kinase complex (*Kamada et al., 2000*). Alternatively, autophagy-related genes could be up-regulated via two transcription factors REPTOR (repressed by TOR) and REPTOR-BP (REPTOR-binding partner) (*Tiebe et al., 2015*). In this study, we identify a novel gene *REPTOR2* in pea aphids by comparing the transcriptome of wingless-destined aphids vs. winged-destined aphids at 24 h post birth. Since *REPTOR2* has a higher expression in the thorax section of wingless-destined aphids, we hypothesized that aphid *REPTOR2* activates autophagic degradation in wing disc to result in the wingless morph. Likewise, winged-destined aphids have a stronger TOR activity to suppress autophagy in wing disc to develop into full wings. Experimentally, we specifically determined: (i) the time point to initiate PCD in the wing disc, (ii) the PCD type that was responsible for wing disc degeneration, and (iii) the role of REPTOR2 and TOR signaling cascade in determination wing dimorphism.

## Results

### Effects of maternal density and duration time on the proportion of winged offspring

To efficiently sample winged offspring, we examined how maternal density and duration time affected the proportion of winged offspring. Compared to the group with a single aphid, groups of 2, 4 and 8 individuals each contacted for 8 h led to higher proportions of winged offspring, respectively (Fig. 1A). Since 2 adult individuals were sufficient to produce over 80% winged offspring, we then tested different contact duration times on the proportion of winged morph (Fig. 1B). Apparently, 4 h of maternal contact duration was sufficient to induce > 90% winged offspring, and 2 h could induced a moderate effect (approximately 50%) (Fig. 1B). Furthermore, group of 2 adult individuals contacting for 4 h could continuously produce over 80% winged offspring for 3 days, followed by a small reduction afterward (Fig. 1C). Furthermore, to eliminate a strain specific induction, other 3 strains of pea aphid were also tested by 2 adults contacting for 4h, and the lowest proportion of winged offspring among 4 aphid strains exceeded 40% (Fig. S4).

**Fig.1.**
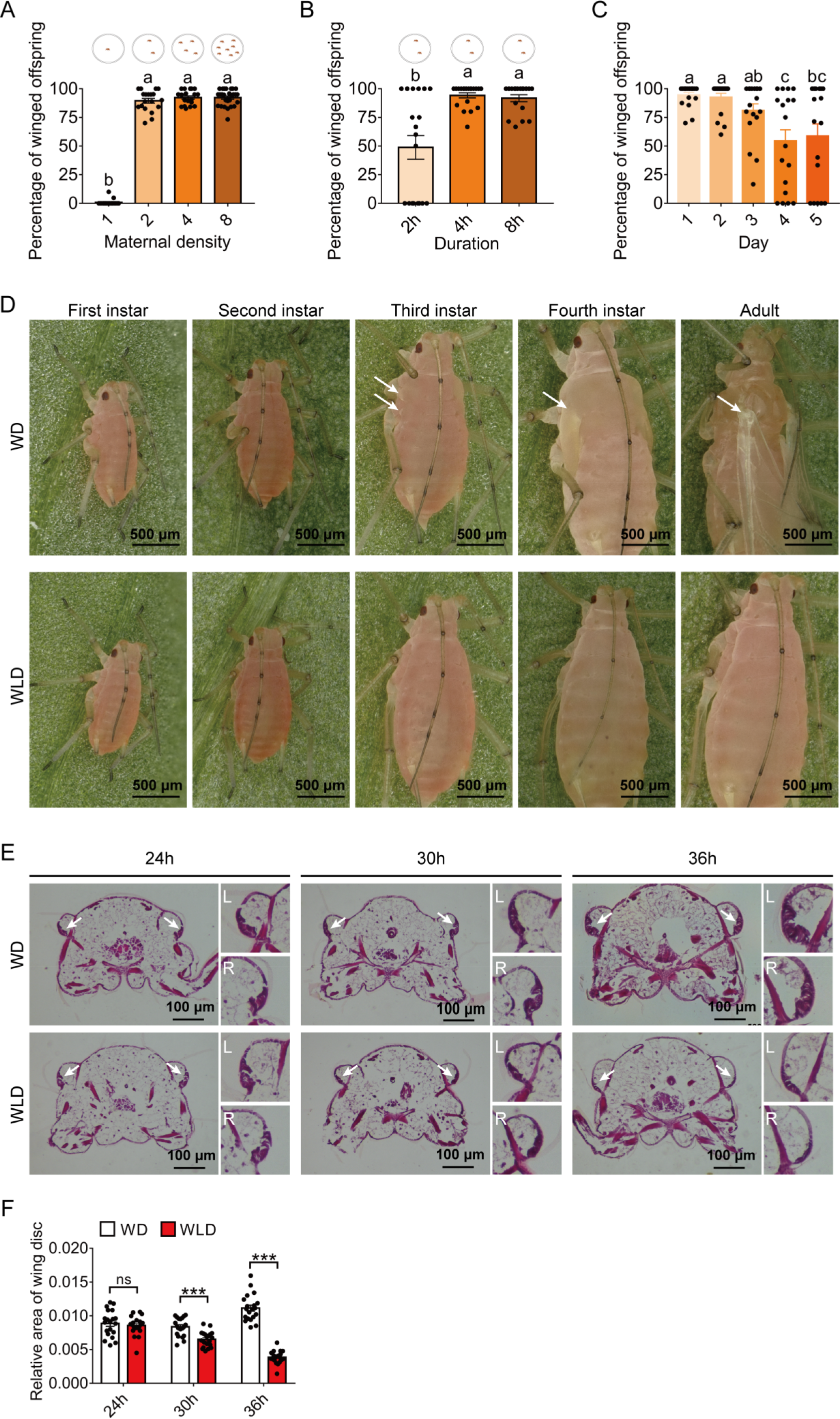
Wing dimorphism in pea aphids is transgenerational with high sensitivity to maternal density and duration time, and 1^st^ instar nymphs at 30 h post birth is a critical stage for developmental plasticity of the wing disc. (A) Two female adults in a petri dish can efficiently induce a high proportion of winged offspring. (B) Percentage of winged offspring produced by the two-adult contacting treatment for different durations. (C) Daily percentage of winged offspring for the two-adult contacting treatment for 4 h. (D) Developmental morphology of aphid wings from the 1^st^ instar nymphal to adult stages in winged and wingless aphids. White arrows: locations of the wing bud or wing in the winged morph. (E) Histological comparisons of wing disc in the 1^st^ instar nymphs at 24 h, 30 h and 36 h post birth, and wing discs existed in both winged- and wingless-destined aphids. White arrows indicate the wing disc. L: left wing disc, R: right wing disc. (F) The relative area of the wing disc measured by imageJ. WD: winged-destined, WLD: wingless-destined. Values in bar plots represent means (±SE) of at least 16 replicates. Tukey’s multiple range tests at *P*<0.05 were used to compare means, and different lowercase letters indicate significant differences. Significant difference at *P*<0.001 is indicated by asterisks (***).

### Developmental differentiation in the wing and wing disc of winged vs. wingless morphs

Previous studies indicated that the wing disc was degenerated at the 2^nd^ instar in wingless morph of the pea aphid (*Ishikawa et al., 2008; Ogawa and Miura, 2013*). We compared the wing morphology across different instar nymphs between wing morphs. Winged- and wingless-destined aphids could not be distinguished from wing morphogenesis until the 3^rd^ instar nymphal stage (Fig. 1D). The wing bud of winged-destined aphids became visible during the 3^rd^ and 4^th^ instar, and developed into full unfolded wing once adult emergence. By contrast, wingless-destined aphids did not possess obvious wing bud at the 3^rd^ and 4^th^ instar stages. To further determine the time point of initiation of the wing disc degeneration, the 1^st^ instar nymphs at 24 h, 30 h, and 36 h post birth were collected for the histological analysis. The wing disc sizes were not significantly different 24 h post birth between the two morphs, but were then rapidly degenerated in wingless-destined morphs while continuously developed and expended in winged-destined morphs 30 h post birth (Fig. 1E and F).

### Autophagic rather than apoptotic degradation was responsible for degeneration of wing disc

Since wing disc degeneration initiating at 30 h post birth, PCD types and processes occurred in wing disc at this time point were characterized by transmission electron microscopy (TEM), immunofluorescence, TUNEL and qPCR. Numerous cytoplasmic vacuoles, large autophagosomes sequestering visible remnants of organelles, and membranous whorls were observed by TEM within the cells of the wing disc of wingless-destined morphs, suggesting typical autophagic features. By contrast, only mitophagy and a few cytoplasmic vacuoles were found in wing disc cells of winged-destined morph. Signs of autophagy were observed in wing disc of 54% cells of the wingless-destined morph, while only 15% cells with the sign of autophagy were found in winged-destined aphids (Fig. 2A). Furthermore, typical features of apoptosis including cell shrinkage, plasma membrane blebbing, apoptotic body, chromatin condensation, nucleolus disorganization, and nuclear fragmentation were barely observed from TEM in wing discs of both winged- and wingless-destined morphs.

**Fig.2.**
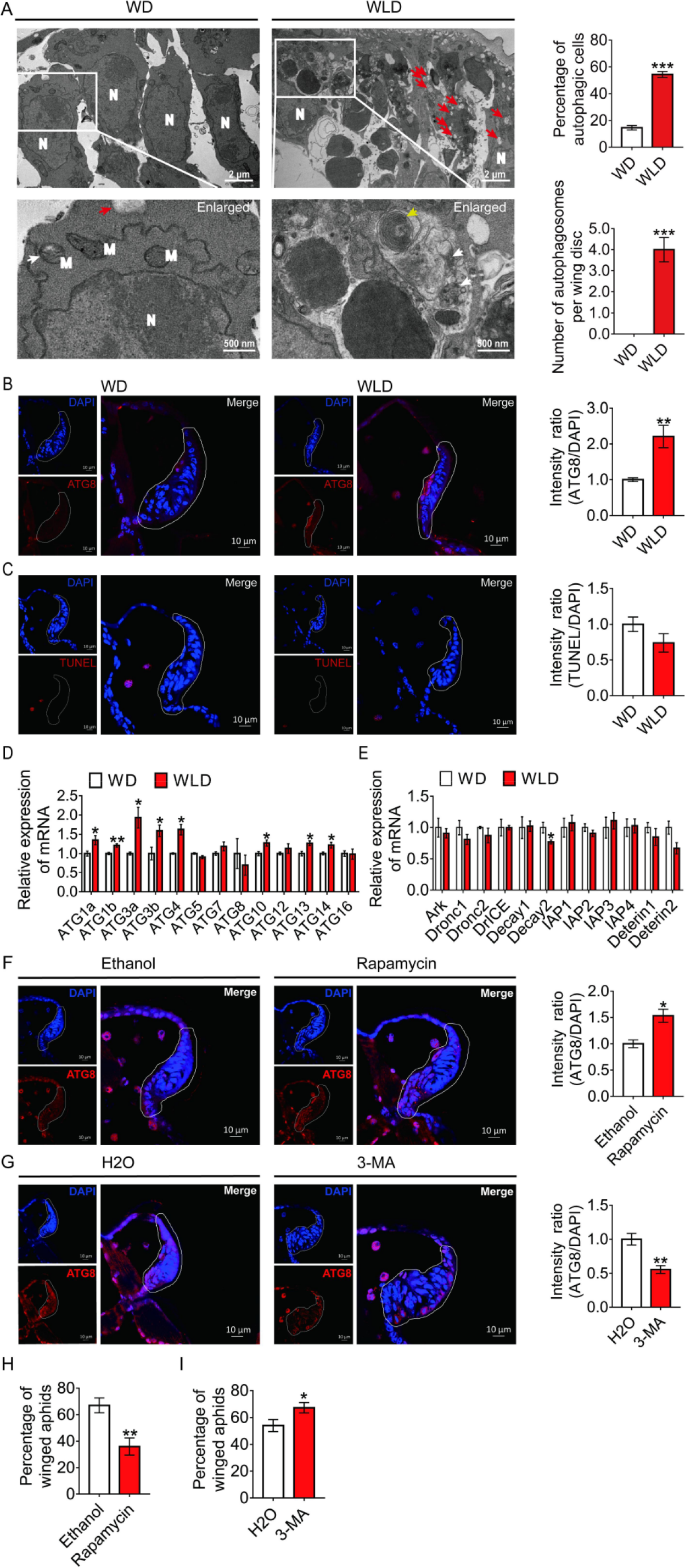
Autophagy, and not apoptosis, is responsible for the degeneration of the wing disc in wingless-destined aphids. (A) TEM images of wing discs of winged- and wingless-destined aphids at 30 h post birth. N: nucleus, M: mitochondrion. Yellow and white arrows indicate autophagosomes containing membranous whorls and remnants of cellular organelles, respectively. Red arrows indicate the vacuoles (n=3). (B) Autophagy in the wing disc was indicated by the hallmark ATG8 (red) in immunofluorescence. The nuclei (blue) were stained by DAPI (n=7). (C) Apoptosis was determined by TUNEL assays (red) (n=5). (D) Autophagy-related genes (*ATG1a*, *ATG1b*, *ATG3a*, *ATG3b*, *ATG4*, *ATG5*, *ATG7*, *ATG8*, *ATG10*, *ATG12*, *ATG13*, *ATG14*) were determined by qPCR (n=4). (E) Pro-apoptotic genes (*Ark*, *Dronc1*, *Dronc2*, *DrICE*, *Deterin1*, *Deterin2*), anti-apoptotic genes (*IAP1*, *IAP2*, *IAP3*, *IAP4*, *Decay1*, *Decay2*) were determined by qPCR (n=4). (F) The effect of autophagy agonist rapamycin, and (G) autophagy inhibitor 3-MA on autophagy in the wing disc. Autophagy was noted by ATG8 (red) in immunofluorescence, and nuclei were stained by DAPI (blue). All relative intensity was quantified by ImageJ (n=5). (H) The effect of autophagy agonist rapamycin, and (I) autophagy inhibitor 3-MA on the proportion of winged aphids. Ethanol and H_2_O were used as control, respectively (n=9). Wing disc shown in confocal was highlighted by white circle. WD: winged-destined, WLD: wingless-destined. Values in bar plots are means (±SE). Independent sample *t*-test was used to compare means, and significant differences between treatments are indicated by asterisks, \**P*<0.05, \*\**P*<0.01, \*\*\**P*<0.001.

The immunofluorescence assay and TUNEL results showed that autophagy, instead of apoptosis, was activated in wing disc cells of the wingless-destined morph (Fig. 2B, 2C). The autophagy-related genes *ATG1a*, *ATG1b*, *ATG3a*, *ATG3b*, *ATG4*, *ATG10*, *ATG13*, and *ATG14* had higher expression in the wingless-destined morph (Fig. 2D), while all apoptosis-related genes except for *Decay2* were not significant between two wing morphs (Fig. 2E). These results indicated that autophagy rather than apoptosis was responsible for the degeneration of the wing disc in pea aphids, and thereby reduced the proportion of the winged morph. To verify this hypothesis, the group with 2 adult individuals within a petri dish for 2 h was applied to produce a moderate proportion (approximately 50%) of winged offspring (Fig. S1). Newborn nymphs were fed with autophagy agonist rapamycin or antagonist 3-MA for 24 h. As anticipated, rapamycin enhanced autophagy in the wing disc of 1^st^ instar nymphs and resulted in a decline of winged proportion from 67% to 36% (Fig. 2F and 2H), while 3-MA attenuated autophagy in the wing disc and increased the winged aphids from 54% to 67% (Fig. 2G and 2I).

### A novel gene *REPTOR2* in pea aphid positively regulated the wing disc autophagy

The 1^st^ instar nymph of wingless-destined morphs had 387 differentially expressed genes in total, with 220 higher expressed genes and 167 lower expressed genes relative to winged-destined morphs at 24 h post birth (Fig. 3A and Dataset S1). Strong evidence has suggested that autophagic cell death could be regulated by TORC1 (*Berry and Baehrecke, 2007; Denton et al., 2012*), we therefore focused on the TOR signaling pathway (Fig. 3B and Dataset S2). Only *REPTOR2* (*repress by TOR 2*, LOC103309809) and *SLC38A9* (*sodium-coupled neutral amino acid transporter 9*, LOC100158916) in the TOR pathway had significantly higher expression in wingless-destined morphs. *SLC38A9* is a lysosomal membrane-resident protein competent in the amino acid transport, which is responsible for amino acids-induced TORC1 activation (*Rebsamen et al., 2015*). *REPTOR* is recognized as a transcription factor that relays TOR signaling cascade by activating the expression of autophagy-related genes (*Tiebe et al., 2015*).

**Fig.3.**
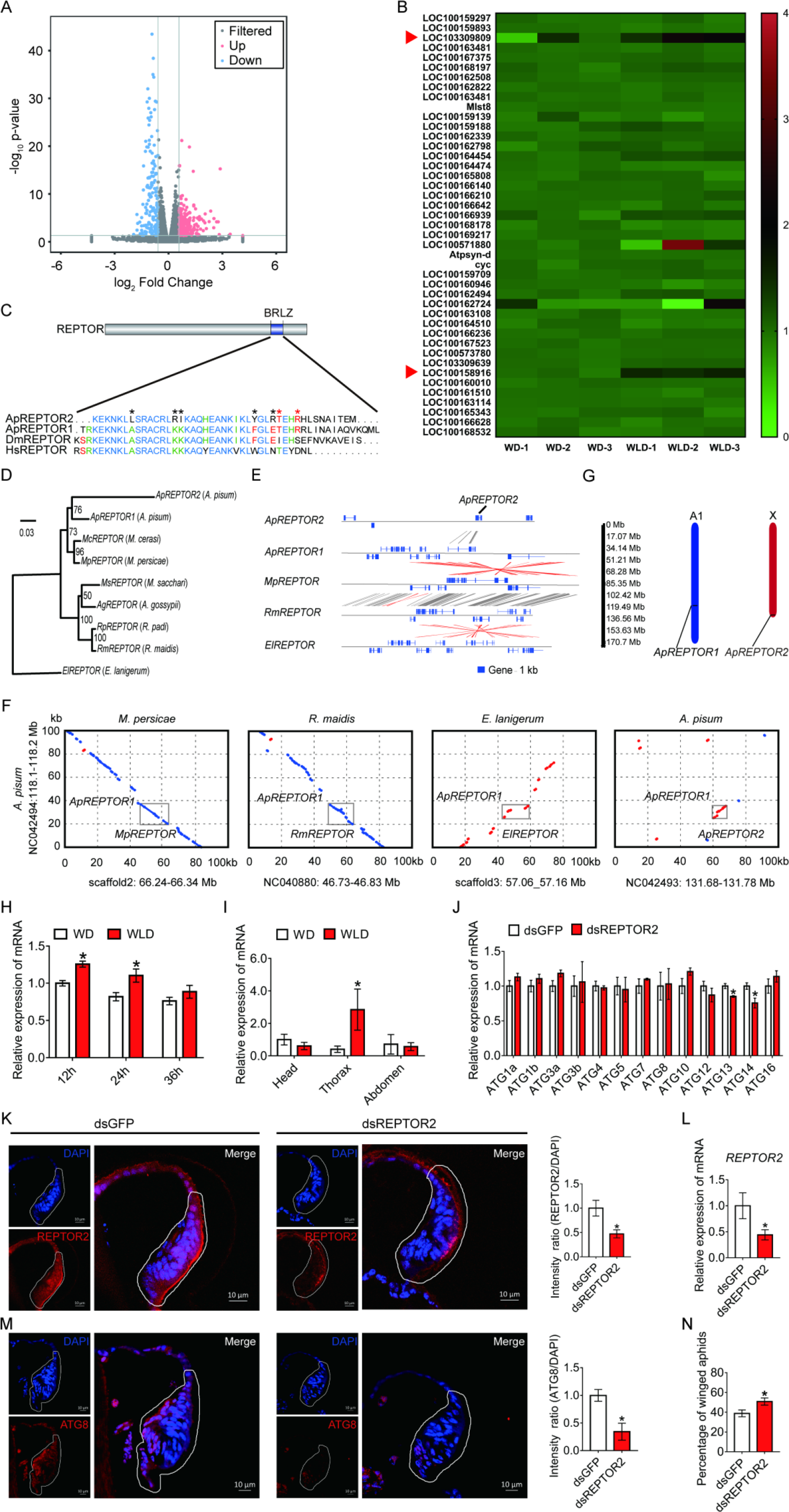
The pea aphid-specific gene *REPTOR2* is highly expressed in wingless-destined morph that activates autophagy in wing disc. (A) The volcano plot of log-transformed FPKM showing the number of differentially expressed genes (DEGs) in wingless- vs. winged-destined aphids at 24 h post birth. Three biological replications were conducted in the RNA-seq analysis. (B) Transcripts mapping to TOR signaling pathway of pea aphids and DEGs in two wing morphs. Red arrows indicated two DEGs, *REPTOR2* (LOC103309809) and *sodium-coupled neutral amino acid transporter 9* (LOC100158916), in wingless- vs. winged-destined aphids. (C) Homology analysis of pea aphid REPTOR2. The amino acid sequence of conserved basic region leucine zippers (BRLZ) domain of pea aphid REPTOR2 was aligned with pea aphid REPTOR1 (ApREPTOR1), human REPTOR (HsREPTOR) and Drosophila REPTOR (DmREPTOR). Red and black asterisks indicated pea aphid-specific variation and ApREPTOR2-specific variation in amino acid, respectively. (D) An unrooted phylogenetic tree was constructed by the IQ-TREE method with coding sequence alignments for aphid REPTOR. The sequences of REPTOR identified from 8 aphid species as well as the candidate pea aphid *REPTOR2* were compared. (E) Sequence comparisons of the LOC103309809 (*ApREPTOR2*) locus and its homologous regions in aphids. Homologous sequences were linked by grey dashes corresponding to the positive strand and by red dashes corresponding to the negative strand. (F) Grey boxes highlight the similarity of aphid REPTOR across 100 kb covering the REPTOR loci. (G) Chromosomal distribution of two REPTOR genes in pea aphids. (H) Relative expression levels of *REPTOR2* in 1^st^ instar nymph at 12 h, 24 h and 36 h post birth in winged- and wingless-destined aphids (n=4). (I) Expression levels of *REPTOR2* in head, thorax and abdomen of winged- and wingless-destined aphids at 24 h post birth (n=4). (J) The effects of ds*REPTOR2* on the expression of genes related to autophagy in whole body of the 1^st^ instar nymphs (n=4). (K) Knockdown of *REPTOR2* reduced its expression in wing disc at 24 h post birth. Fluorescence in situ hybridization (FISH) was used to detect the *REPTOR2* expression level in aphid wing discs. *REPTOR2* was hybridized with 5-CY3 in red, and nuclei were stained with DAPI in blue (n=4). (L) *dsREPTOR2* feeding decreased the gene expression of *REPTOR2*, as determined by qPCR (n=4). (M) Knockdown of *REPTOR2* resulted in the decline of autophagy in the wing disc at 30 h post birth. Autophagy was noted by ATG8 (red) in immunofluorescence, and nuclei were stained by DAPI (blue) (n=4). (N) *dsREPTOR2* treatment increased the proportion of winged aphids (n=10). Wing discs shown in confocal were highlighted by white circles. WD: winged-destined, WLD: wingless-destined. Values in bar plots are means (±SE). Independent sample *t*-test was used to compare mean, and significant differences between treatments are indicated by asterisks: \**P*<0.05, \*\**P*<0.01, \*\*\**P*<0.001.

When aligned with *Acyrthosiphon pisum* genome, two REPTOR homologs, *ApREPTOR1* (LOC100168197) and *ApREPTOR2* (LOC103309809), were identified. Seven other aphid species contained only a single REPTOR copy, including *Myzus cerasi*, *Myzus persicae*, *Melanaphis secchari*, *Aphis gossyppii*, *Rhopalosiphum padi*, *Rhopalosiphum maidis*, and *Eriosoma lanigerum*. Both *REPTOR* in *A. pisum* consisted of a conserved basic region leucine zipper (BRLZ) domain (Fig. 3C). An unrooted phylogenetic tree with bootstrap consensus was constructed across 8 species of aphid using coding sequence alignments of *REPTOR* (Figure 3D). MUMmer analyses were used to determine the similarity and homologous regions of *ApREPTOR2* in 4 aphid species with chromosome-level genomes in Aphidbase. *ApREPTOR1* and *MpREPTOR* had 9 exons, while *ApREPTOR2* had 7 exons (Fig. 3E). *ApREPTOR1* shared high sequence similarity with *MpREPTOR*, *RmREPTOR*, and *ApREPTOR2*, but not with *ElREPTOR*. The variations in the flanking region of REPTOR indicated that *ApREPTOR2* was quite different from *ApREPTOR1* (Fig. 3F). Chromosomal distribution results further showed that *REPTOR1* was located in A1 chromosome while *REPTOR2* was located in X chromosome of *A. pisum* (Fig. 3G). These results indicated that *REPTOR1* in pea aphid may have experienced a gene duplication to generate a novel gene *REPTOR2* which probably translocate from autosome A1 to the sex chromosome.

*REPTOR2* had higher expression in wingless-destined morphs at 12 h and 24 h post birth than that of winged-destined morphs (Fig. 3H), and highly expressed in the thorax of wingless-destined morphs (Fig. 3I), suggesting that *REPTOR2* was essential in activating autophagic degradation of the wing disc. Furthermore, newborn nymphs fed with *dsREPTOR2-RNA* reduced the *REPTOR2* transcripts in the wing disc and whole body by 53% and 56%, respectively (Fig. 3K and 3L), and decreased gene expressions of *ATG13* and *ATG14* (Fig. 3J). Knockdown of *REPTOR2* attenuated autophagy in the wing disc (Fig. 3M), resulting in higher proportion of the winged morph, from 39% to 51% (Fig. 3N).

### The effect of TOR on wing disc autophagy and the proportion of the winged morph

Suppression of TOR plays a causative role in inducing autophagic cell death during tissue degeneration in insect (*Berry and Baehrecke, 2007; Denton et al., 2012*). As expected in wing disc, the fluorescence in situ hybridization (FISH) data indicated that *TOR* had higher abundance in winged- than wingless-destined morphs (Fig. 4A). Feeding newborn nymphs with *dsTOR-RNA* reduced the *TOR* transcripts in whole body and wing disc by 48% and 34%, respectively (Fig. 4B and 4C), which led to decline of winged proportion from 69% to 52% (Fig. 4D). By contrast, activation of TOR via pharmacological agonist MHY1485 suppressed autophagy in wing disc (Fig. 4E), and increased winged proportion from 37% to 55% (Fig. 4F). Furthermore, suppression of TOR via depletion of amino acids by 50% enhanced autophagy in wing disc (Fig. 4G), and decreased winged proportion from 49% to 15% (Fig. 4H).

**Fig.4.**
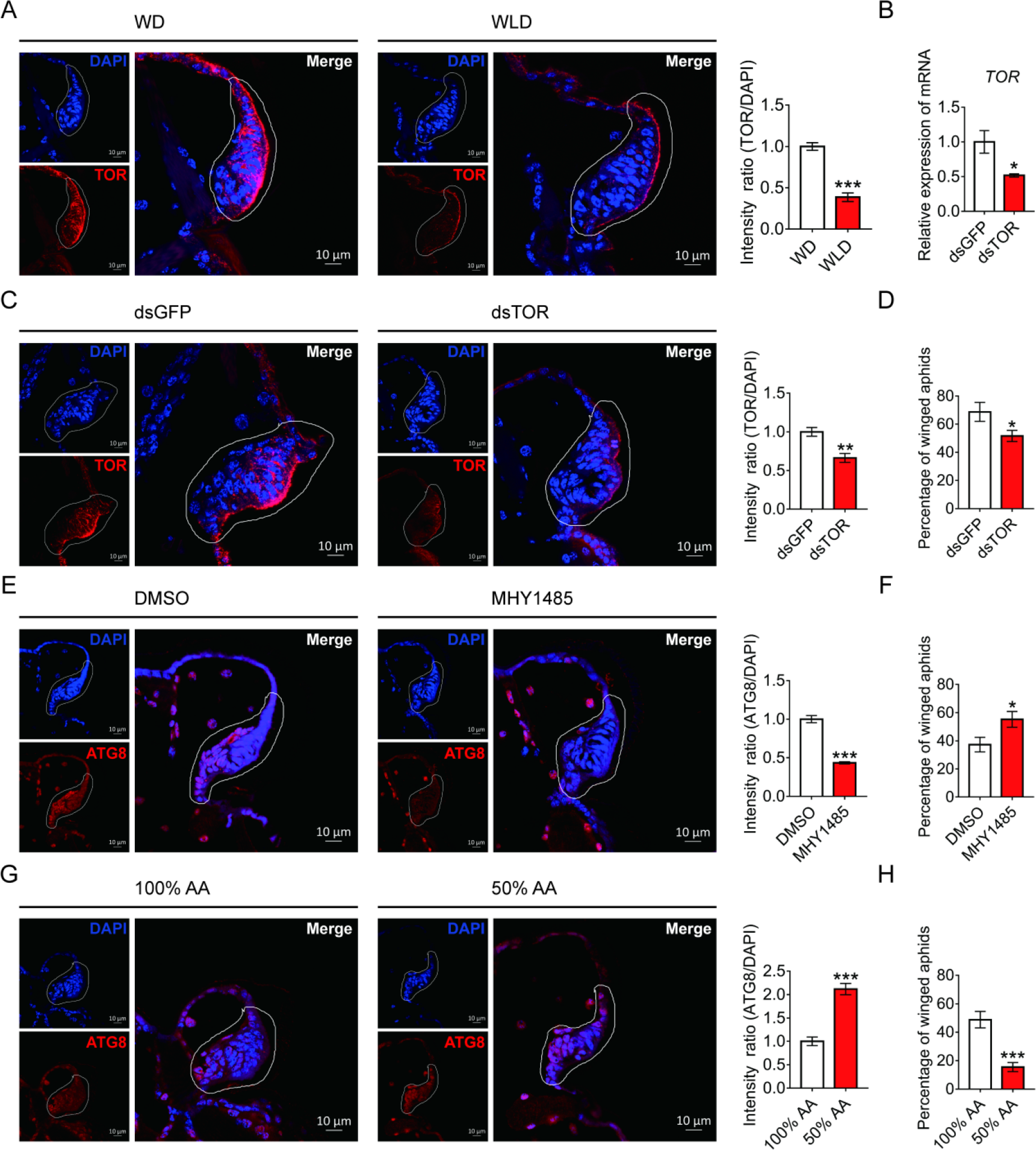
TOR negatively regulated autophagy in the wing disc and positively affected the proportion of winged aphids. (A) *TOR* expression in the wing disc of 1^st^ instar nymph of winged- and wingless-destined aphids at 24 h post birth, as determined by mRNA-FISH. *TOR* was hybridized with 5-CY3 in red, and nuclei were stained with DAPI in blue (n=9). (B) Newborn nymphs fed with dsRNA reduced *TOR* expression in whole body 24 h post birth, as determined by qPCR (n=4). (C) Knockdown of *TOR* reduced its expression in the wing disc 24 h post birth, shown in mRNA-FISH (n=4), and (D) increased the proportion of winged aphids (n=10). (E) Application of TOR agonist MHY1485 attenuated autophagy in the wing disc 30 h post birth (n=9). (F) Activation of TOR increased the proportion of winged aphids (n=8). (G) Amino acids supplied at 50% of the standard diet (50% AA) activated autophagy in the wing disc 30 h post birth (n=8). (H) 50% AA reduced the proportion of winged aphids (n=8). Autophagy in the wing disc was indicated by the hallmark ATG8 (red) in immunofluorescence. The nuclei (blue) were stained by DAPI. Relative intensity was quantified by ImageJ. Wing discs shown in confocal was highlighted by white circles. WD: winged-destined, WLD: wingless-destined. Values in bar plots are means (±SE). Independent sample *t*-test was used to compare mean, and significant differences between treatments are indicated by asterisks: \**P*<0.05, \*\**P*<0.01, \*\*\**P*<0.001.

## Discussion

The wing disc consists of undifferentiated and proliferating cells designated to develop into a full insect wing during the nymphal stage (*Neto-Silva et al., 2009*). Fate of the wing disc is possibly determined by maternal signals triggered by environmental challenges such as density, physical contact and natural enemy, as wing dimorphism in aphids is transgenerational (*Braendle et al., 2006*). It has been reported that ecdysteroids and miR-9b are the molecules that transduce maternal crowding signals to embryo that leads to a high proportion of winged offspring (*Vellichirammal et al., 2017; Shang et al., 2020*), but relative few studies have focused on the role of the wing disc degradation at the onset of the nymphal stage in forming wing dimorphism. Our study demonstrated that a novel gene REPTOR2 in pea aphid had high expression in the 1^st^ instar nymphs of wingless-destined morph 12 h post birth, which activated autophagy in the wing disc and resulted in its degeneration as early as 30 h post birth. Different types of PCD has been reported to activate in tissue degeneration in many insect orders (*Xu et al., 2019*). Wing discs of soldier ant and some sex-dependent wingless morphs of lepidopteran insects are degenerated by apoptotic cell death (*Fujiwara and Ogai, 2001; Niitsu, 2001; Sameshima et al., 2004; Rajakumar et al., 2012; Rajakumar et al., 2018*). On the other hand, apoptotic cell death did not seem to be necessary in wing disc degradation in aphids. Instead, typical features of autophagic cell death occurred at 30 h post birth in wingless-destined morphs. Our results support a model concerning aphid wing disc degeneration and development: maternal aphids experience crowding could increase TOR in 1^st^ instar nymphs. This is followed by the repression of REPTOR2 and autophagic degradation in the wing disc, thereby causing a high proportion of winged offspring. For the solitary maternal group, TOR activity is arrested in the 1^st^ instar offspring, and REPTOR2 and autophagy are subsequently activated in wing disc, leading to a high proportion of wingless offspring (Fig. 5). Given high expression in thorax of wingless-destined morph at early 1^st^ instar nymphal stage, it was likely that REPTOR2 amplified the TOR signaling to activate autophagic degeneration of the wing disc in a tissue-specific manner.

**Fig. 5.**
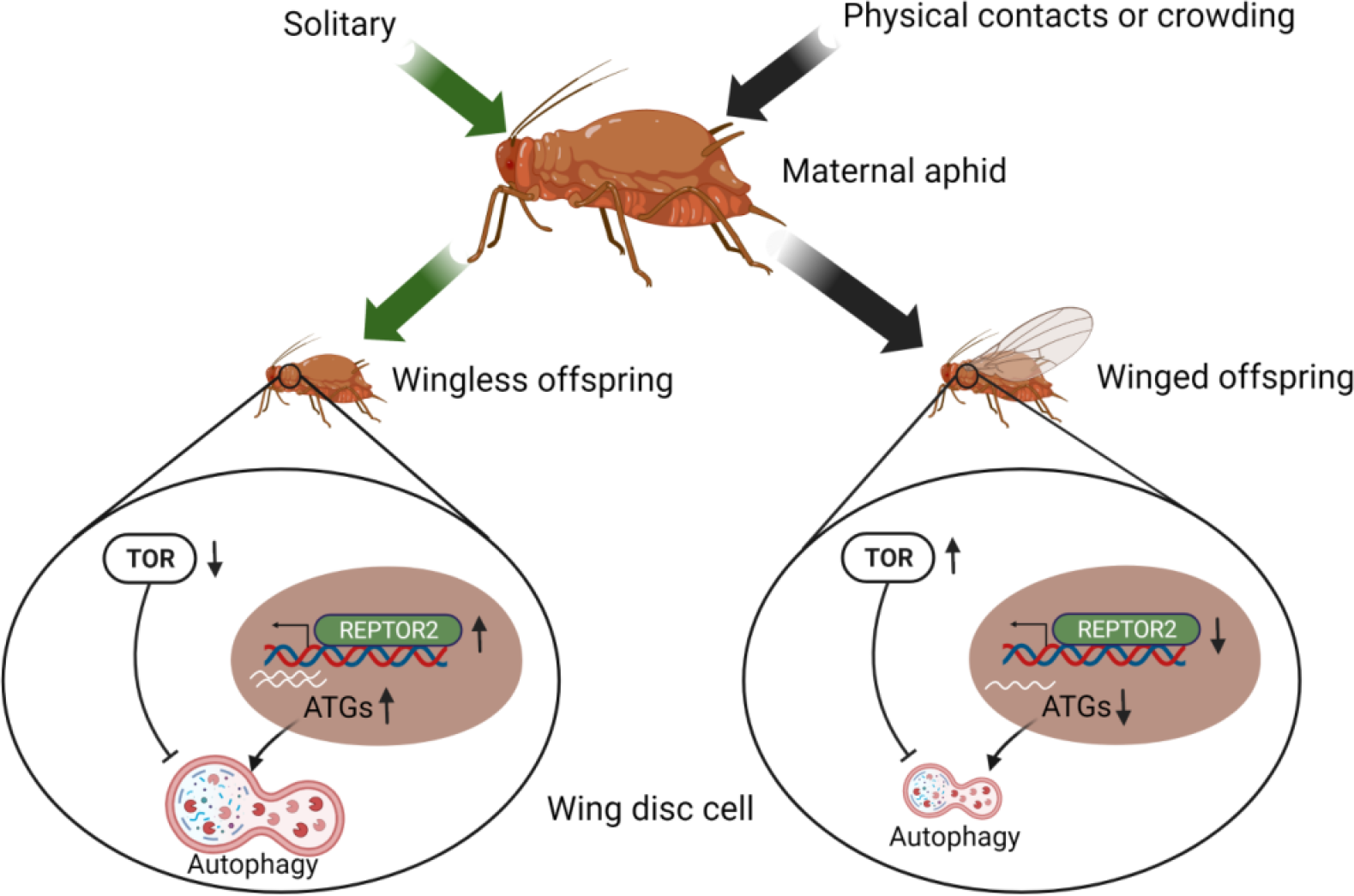
TOR signaling pathway-controlled wing disc degeneration of the 1st instar of pea aphid nymphs. All treatments, indicated along the top, were applied to wingless maternal aphids. Arrows correspond to the treatments, coded by different colors. TOR: target of rapamycin, REPTOR2: repress by TOR 2, ATGs: autophagy related genes. This graph was created with BioRender.com.

The TOR signaling pathway has been reported to modulate developmental plasticity in some insects (*Koyama et al., 2013*). For instance, activation of TOR activity in honeybees is responsible for the developmental transition from facultatively sterile workers to reproductive queens, while repressing TOR activity could induce worker characters in queen-destined individuals (*Michalak et al., 2007*). Transcriptomic analysis in maternal crowding of pea aphids suggested that genes functionally related to TOR signaling pathway were differentially expressed in mother aphids (*Parker et al., 2021*). We therefore hypothesized that the canonical pathway of TOR-REPTOR-autophagy may be responsible for developmental plasticity of the wing disc in aphids, in accordance with lower TOR abundance in wingless-destined morph. Suppression of TOR in Drosophila could up-regulate the expression of autophagy-related genes via phosphorylation of Ser527 and Ser530 in REPTOR (*Tiebe et al., 2015*). Similarly, pea aphid *REPTOR2* is predicted to contain a conserved basic region leucine zipper (BRLZ) domain which is capable of binding to GTAAACAA, the binding motif of FOXO as well (*Tiebe et al., 2015*). FOXO is a transcription factor that relays the insulin signaling to insect wing dimorphism. In the planthopper, activating insulin signaling transforms wing commitment from short-winged to long-winged morph via repressing the FOXO activity (*Xu et al., 2015*). Activating FOXO activity inhibits development of long-winged morph by suppressing vestigial (Vg) expression during the wing-morph decision stage (*Zhang et al., 2021*). It appears that FOXO and REPTOR2 respectively relay the insulin and TOR signaling cascades to control wing dimorphism in planthoppers and aphids. Alternatively, the BRLZ domain is required to form REPTOR/REPTOR-BP homo/hetero-dimer complex which conveys negative regulation of TOR signaling on genes related to autophagy (*Tiebe et al., 2015*). Our results are consistent with the canonical pathway regulated by TOR that either knockdown of *REPTOR2* or activation of TOR could attenuate autophagy in wing disc. These results suggest that *REPTOR2* from pea aphids served as a transcription factor to exert a negative effect of TOR signaling on autophagy initiation.

Insulin signaling is known to determine the wing-morph switch in planthoppers (*Xu et al., 2015*), and is also responsible for production of winged offspring regulated by miR-9b-ABCG4 cascade in brown citrus aphid (*Shang et al., 2020*). Furthermore, insulin could enhance TOR signaling by releasing the negative feedback of Akt on the tuberous sclerosis proteins TSC1 (hamartin) and TSC2 (tuberin) (*Blagosklonny et al., 2009*). It has been shown that embryos derived from pea aphids subjected to crowding treatment have higher insulin signaling (*Grantham et al., 2019*). This may lead to stronger TOR signaling in the 1^st^ instar offspring, which could suppress wing disc autophagy and increase the proportion of winged aphids.

Gene duplication is a major source for novel gene production (*Kaessmann, 2010; Prince and Pickett, 2002*), which has become an important mechanism regulating phenotypic plasticity (*Chen et al., 2013*), including manipulating wing polyphenism in insects. For instance, a derived insertion of a duplicated *fs3* (follistatin 3) determined the wingless morph formation for male pea aphids (*Li et al., 2020*). Another duplicated gene *InR2* (insulin receptor 2) was highly expressed in the wing bud of the planthopper that served as a switch regulator of short-winged morph formation (*Xu et al., 2015*). Similarly, different from other 7 aphid species with single copies of REPTOR, pea aphids possess two REPTOR genes. Although it has lost 2 exons, *ApREPTOR2* has higher similarity in coding sequence with *ApREPTOR1*, but shares little similarity in the flanking regions. This is in agreement with the chromosomal distribution results indicating that *ApREPTOR1* is located in A1 chromosome while *ApREPTOR2* is located in X chromosome. *ApREPTOR2* is likely a duplicate from *ApREPTOR1*, followed by translocating from A1 to X chromosome. This may also warrant reproductive role of sexual female in pea aphid, because most sexual females are monomorphic wingless, whereas males that contain just single X chromosome are dimorphic (*Brisson, 2010*).

Numbers of diverged paralogs in pea aphid could be differentially expressed in response to particular environmental conditions by which extensive polyphenisms are exhibited (*Shigenobu et al., 2010; Brisson et al., 2010*). Although *ApREPTOR2* is significant for the winged- vs. wingless-destined aphids and controls the wing disc degradation, the expression of *ApREPTOR1* was unaffected between wing morphs in head, thorax, and abdomen at 24 h post birth (Fig. S3A). Knockdown of *ApREPTOR1* did not affect the expression of *ApREPTOR2* and *ATG*s and the proportion of winged aphids (Fig. S3B-E). These results indicated that *ApREPTOR2* rather than the canonical *REPTOR* could activate a tissue-specific degradation in wing disc by preventing a systemic PCD. Apparently, more research is necessary to further elucidate how other aphid species that contain single *REPTOR* relay the TOR signaling on autophagic degradation in the wing disc.

## Materials and methods

### Aphid rearing

The pea aphid *Acyrthosiphon pisum* (strain: Ningxia Red) was originally collected from *Medicago sativa* in Ningxia Province and had been reared in the laboratory for over 8 years. Nymphs from the same parthenogenetic pea aphid female were reared on *Vicia faba* at 18-20 °C, with 60% relative humidity and a photoperiod of a 16 h: 8 h (light/dark) cycle. To eliminate the transgenerational effects on offspring morphs, females were maintained at low density (three per plant) on *V. faba* seedlings for more than three generations (*Sutherland, 1969*). Other 3 strains of *A. pisum*, including Ningxia Green, Gansu Red and Gansu Green, were originally collected from *Medicago sativa* in Ningxia Province and Gansu Province respectively, and were used to test the inductive effect of two-adult contacts on the proportion of winged offspring.

### Induction of winged- and wingless-destined morphs

In *A. pisum*, physical contact is known to be the key stimulus inducing the production of winged viviparous females (*Sutherland, 1969*). To effectively induce high proportion of winged offspring, the maternal density and contacting duration were evaluate within petri dishes (35 mm in diameter). For maternal density, 4 groups consisted of 1, 2, 4, or 8 wingless adults respectively, were placed in a petri dish for 8 h. For the contacting duration, two wingless adults were placed in a petri dish for three different durations: 2 h, 4 h, and 8 h. After treatment, each adult was then transferred to fresh detached leaf of *V. faba* that kept in petri dishes with 1% agar, and allowed to reproduce for 24 h. After newly born nymphs reached the 4^th^ instar, the proportion of winged offspring from each adult was calculated. Furthermore, 2 wingless adults were placed in petri dish for 4 h. Adults were then transferred to *V. faba* plants, one female per plant, to oviposit for five days and adults were discarded. Offspring produced by each day were transferred to detached leaf of *V. faba* that kept in petri dishes with 1% agar. Fresh leaves were changed every three days. Winged or wingless of each offspring was counted until the 4^th^ instar. To eliminate the strain specific induction, aphid strains of Ningxia Green, Gansu Red, and Gansu Green were also tested using 2 wingless adults contacting for 4h.

### Developmental time and wing morphology of winged- and wingless-destined morphs

Newly born nymphs of winged- or wingless-destined morphs were raised individually on fresh, detached leaf of *V. faba* in petri dishes. The nymphal developmental time and the pre-reproductive period were recorded every 12 h. The digital microscope (Keyence VHX-5000, Osaka, Japan) was used to determine the wing morphology in nymphs of each instar.

### Histology

Winged- and wingless-destined morphs at 24 h, 30 h, and 36 h post birth were collected and placed in 4% paraformaldehyde fixative overnight at 4 °C, washed in PBS for three times and dehydrated at 4 °C for 24 h in 30% sucrose solution. After freezing at −20°C in Tissue Freezing Medium (Leica), aphids were sectioned at 10 μm using a Leica CM1950 freezing microtome, and stained with H & E (Beyotime, C0105) according to the standard protocol. Images of wing discs were taken using a Nikon light microscope, equipped with a DS-Fi1c camera (Nikon), and the images were generated using the NIS-Element D software (Nikon). Relative area of wing disc was analyzed using ImageJ software.

### Transmission electron microscope

Since wing disc degeneration initiated at 30 h post birth, winged- and wingless-destined morphs at this time point were collected and fixed with 2.5% (vol/vol) glutaraldehyde, and washed four times in Phosphate Buffer (PB) (0.1 M, pH 7.4). Aphids were then fixed with 1% (wt/vol) osmium tetraoxide in PB for 2 h at 4 °C, and dehydrated through a graded ethanol series (30%, 50%, 70%, 80%, 90%, 100%, 100%, 7min each) into pure acetone (2×10min). Samples were infiltrated in a graded mixture (3:1, 1:1, 1:3) of acetone and SPI-PON812 resin (16.2 g SPI-PON812, 10 g dodecyl succinic anhydride (DDSA) and 8.9 g N-methylolacrylamide (NMA), then in pure resin. Aphids were embedded in pure resin with 1.5% benzyl dimethyl amine (BDMA) and polymerized for 12 h at 45°C, and 48 h at 60°C. The ultrathin sections (70nm thick) were sectioned with microtome (Leica EM UC6, Austria), and then double-stained by uranyl acetate and lead citrate, and imaged under transmission electron microscope (FEI Tecnai Spirit120kV, USA). Ultrathin sections were used to count the cell number that had signs of autophagy (mitophagy, cytoplasmic vacuoles, large autophagosomes sequestering visible remnants of organelles and membranous whorls) (*Simonet et al., 2018*). The percentage of autophagic cells was determined by randomly counted cells in the wing disc.

### Immunofluorescence detection and TUNEL assays

The frozen sections of aphids with different treatments were rinsed for three times in TBST (TBS with 0.05% Tween-20). TUNEL assays were carried out by One Step TUNEL Apoptosis Assay Kit (Beyotime, C1089) according to the manufacturer’s protocol. For immunofluorescence, frozen sections of aphids were blocked with SuperBlock T20 (Pierce) for 20min. The samples were incubated with the primary antibody (anti-ATG8, 1:200) overnight at 4°C, rinsed three times in TBST, and incubated with the secondary antibody (1:500) at room temperature for 2 h. A negative control was performed in each independent experiment. The samples were rinsed for three times in TBST, and mounted in Fluoroshield Mounting Medium with DAPI (Abcam). Sections were imaged using a Zeiss LSM710 confocal microscope (Zeiss, Germany). The polyclonal antibody rabbit anti-ATG8 was kindly provided by Professor Le Kang (Institute of Zoology, Chinese Academy of Sciences). Anti-rabbit IgG (H+L), F(ab’)2 Fragment (Alexa Fluor® 594 Conjugate, 8889S) was purchased from CST. Relative intensities of TUNEL and ATG8 were analyzed using the ImageJ software.

### Fluorescence*in situ* hybridization (FISH)

FISH was performed with a technique modified from Kliot et al. (*Kliot et al., 2014*). After washing by TBST (TBS with 0.2%Triton-X) for 10 min, the frozen sections were rinsed for three times in hybridization buffer (20 mM Tris-HCl, pH 8.0, 0.9 M NaCl, 0.01% [wt/vol] sodium dodecyl sulfate, 30% [vol/vol] formamide) for pre-hybrid (without the probe). The frozen sections of aphids were then hybridized overnight in hybridization buffer containing 10 pmol of the fluorescent RNA probe (conjugated with Cy3). Afterward, the frozen sections were rinsed for three times in TBST, and mounted in Fluoroshield Mounting Medium with DAPI (Abcam). Sections were imaged using a Zeiss LSM710 confocal microscope (Zeiss, Germany).

### Quantitative PCR

Total RNA of aphid samples was extracted by TRIzol™ Reagent (Thermo Fisher) according to the manufacturer’s protocol, and cDNA was synthesized using the FastQuant RT Kit with gDNase (Tiangen). The qPCR reactions were performed with the PowerUp SYBR Green Master Mix (Applied Biosystems). RT-qPCR reactions were carried out on the QuantStudio 12K Flex Real-Time PCR System (ABI, A25742) as followed: 30 s at 95°C; followed by 40 cycles of 10 s at 95°C, 30 s at 60°C; and finally one cycle of 15 s at 95°C, 60 s at 60°C, and 15 s at 95°C. The melting curves were used to determine the specificity of the PCR products. The house-keeping gene *actin* (ACYPI000064) was used as the internal qPCR standard to analyze gene expression (*Simonet et al., 2018*). The relative level of each target gene was standardized by comparing the copy numbers of target mRNA with copy numbers of *actin*. Data were analyzed by the 2^−ΔΔCT^ relative quantification method. Specific primers for each gene were designed from the aphid sequences using Primer Premier 6 software (Table S1). The availability of each pair of primers had been tested in the preliminary experiments. The oligonucleotides are listed in Table S1.

### Pharmacological experiments and RNA interference via feeding

Twenty newborn aphids were placed in cylindrical containers, 20 mm diameter × 20 mm height. Each container was provided with a 100 μl artificial diet (*Haribal and Jander, 2015*). The autophagy inhibitor 3-MA (Selleck, S2767) was dissolved in double distilled water and incorporated into diet at 10 μM. The autophagy agonist rapamycin (Selleck, S1039) was dissolved in ethanol and mixed in diet at a concentration of 10 μM, and an equivalent amount of ethanol was used as a feeding control. The TOR agonist MHY1485 (Sellect, S7811) was dissolved in DMSO and mixed in diet at a concentration of 10 μM, and an equivalent amount of DMSO was used as a feeding control. Diet with 50% amino acids (each amino acid was supplied at 50% of standard formulation) was used to repress TOR activity, and a normal diet was used as a control. All chemicals were administered via 24 h feeding.

The T7 RiboMAX Express RNAi System (Promega, P1700) was used to synthesize double stranded RNA (dsRNA) according to manufacturer’s protocol. dsRNAs, i.e. dsREPTOR1,dsREPTOR2, dsTOR (5μl, 10 μg/μl) or dsGFP control were added to 95μl artificial diet to feed aphids. The interference efficiencies were measured by qPCR.

### RNA-seq and data analysis

To determine differentially expressed genes in winged- and wingless-destined morphs, the whole body of 20 aphids at 24 h post birth was collected for further RNA extraction. RNA purity and quantification were evaluated using the NanoDrop 2000 spectrophotometer (Thermo Scientific, USA). RNA integrity was assessed using the Agilent 2100 Bioanalyzer (Agilent Technologies, Santa Clara, CA, USA). The filtered data were obtained by removing low quality reads from the raw data with HTseq-count software (*Anders et al., 2014*). After quality control, paired-end clean reads were aligned to the reference genome downloaded from NCBI (ftp://ftp.ncbi.nlm.nih.gov/genomes/all/GCF/005/508/785/GCF_005508785.1_pea_aphid_22Mar2018_4r6ur/GCF_005508785.1). FPKM of each gene was calculated using Cufflinks (v2.1.1). Differential expression analysis was performed using the DESeq (2012) R package (*Anders et al., 2012*). log2 |fold change| >0.58, p-value<0.05 was set as the threshold for significantly differential expression. Sequence data have been deposited in the National Center for Biotechnology Information’s Sequence Read Archive, https://www.ncbi.nlm.nih.gov/sra (accession no. PRJNA857589).

### Analysis of REPTOR

Two REPTOR genes were identified in the *A. pisum* reference genome (GCF_005508785.1): LOC100168197 (*REPTOR1*) and LOC103309809 (*REPTOR2*). The REPTOR homologs in the *Myzus cerasi*, *Myzus persicae*, *Melanaphis secchari*, *Aphis gossyppii*, *Rhopalosiphum padi*, *Rhopalosiphum maidis*, and *Eriosoma lanigerum* were identified by blastp using pea aphid REPTOR1. Coding sequences of homologous genes of REPTOR, including LOC100168197 (*REPTOR1*, *A. pisum*), LOC103309809 (*REPTOR2*, *A. pisum*), Mca07599.t1 (*M. cerasi*), g26487.t1 (*M. persicae*), LOC112600553 (*M. secchari*), LOC114124537 (*A. gossyppii*), g1773.t1 (*R. padi*), LOC113558611 (*R. maidis*), and jg23767.t1 (*E. lanigerum*) were obtained from NCBI and AphidBase. Sequences were aligned with MUSCLE v3.8.1551 (*Edgar, 2004*). Phylogenetic analysis was performed in IQ-TREE v2.1.4-beta with HKY+F+G4 model (*Minh et al., 2020*) and the phylogenetic tree was visualized in FigTree v1.4.4 (http://tree.bio.ed.ac.uk/software/figtree/). Genomic sequences of orthologous regions of REPTOR were obtained from *M. persicae* (v2.0), *R. maidis* (v1.0), and *E. lanigerum* (v1.0), which have chromosome-level genome assemblies available at Aphidbase. Sequences were compared with NCBI blast v 2.2.26 and homologous sequences were linked by the gray color when sequences matched to positive strand and by the red color when sequences matched to negative strand. Dot plots were generated using MUMmer v3.23. MapGene2Chrom v2.1 was used to analyze the chromosomal distribution of two pea aphid REPTOR genes (http://mg2c.iask.in/mg2c_v2.1/). Conserved domains of REPTOR were identified by The SMART (http://smart.embl-heidelberg.de/). The amino acid sequences of REPTOR homologs from *A. pisum*, *D. melanogaster*, and *H. sapiens* were downloaded from GenBank and aligned with *ApREPTOR2* using MUSCLE v3.8.1551.

### Statistical Analysis

Statistical analyses were performed with SPSS software (Chicago, IL, USA) and GraphPad software. All data were checked for normality by the Wilk-Shapiro test. Student’s t-tests were used to analyze the relative area of the wing disc, percentage of autophagic cells, the number of autophagosomes per wing disc, intensity ratios of ATG8/ DAPI), TUNEL/ DAPI, REPTOR2/ DAPI, and TOR/ DAPI, percentage of winged aphids, and quantification of gene expression for two-group comparisons. ANOVA was used to analyze the effects of maternal density and contacting duration on the proportion of winged offspring, and proportion of winged offspring in each day for 4 h contacting treatment with two female adults.

## Acknowledgements

We thank our colleagues Prof. Jinfeng Chen for his support in genetic analysis and Prof. Chen-Zhu Wang for his advises in project coordination and manuscript writing. We appreciate TEM sample preparing and analysis support from Can Peng, Center for Biological Imaging, Institute of Biophysics, CAS. This project was supported by the Strategic Priority Research Program of the Chinese Academy of Sciences (No. XDPB16), the National Natural Science Foundation of China (Nos. 31970453 and 31870394), and the State Key Laboratory of Integrated Management of Pest Insects and Rodents (No. IPM2206).

## Author contributions

E.Y., B.D., Z.Q. and Y.Y. performing experiments, H.G., W.C. and Y.M. analyzing data; K.Z-S., F.G. and Y.S. project designing and coordination, manuscript writing.

## Supplementary information

**Fig. S1.**
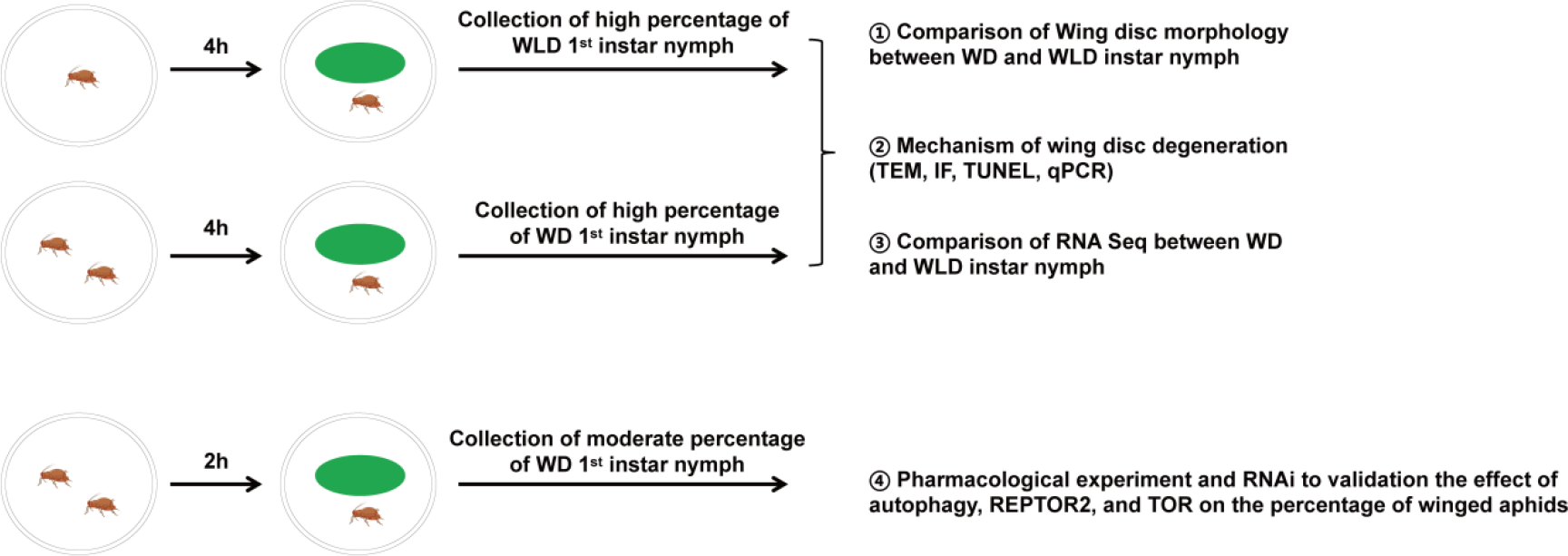
Experimental design and procedure. A flow diagram of the design and sampling procedure for maternal treatment and offspring collection. Female adults of pea aphids were subjected to single adult or two adult individual treatement in the petri dish for 4 h. Since the solitary mother produced high proportion of wingless offspring (>90%), the collected newborn nymphs during the first 2 days named wingless-destined aphids. Likewise, the two-adult individuals contacting treatment for 4 h resulted in high proportion of winged offspring (>90%), so newborn nymphs at first 2 days named winged-destined aphids. Alternatively, two adult individuals contacting for 2 h could cause a moderate proportion of winged offspring (~50%). WD: winged-destined, WLD: wingless-destined.

**Fig. S2.**
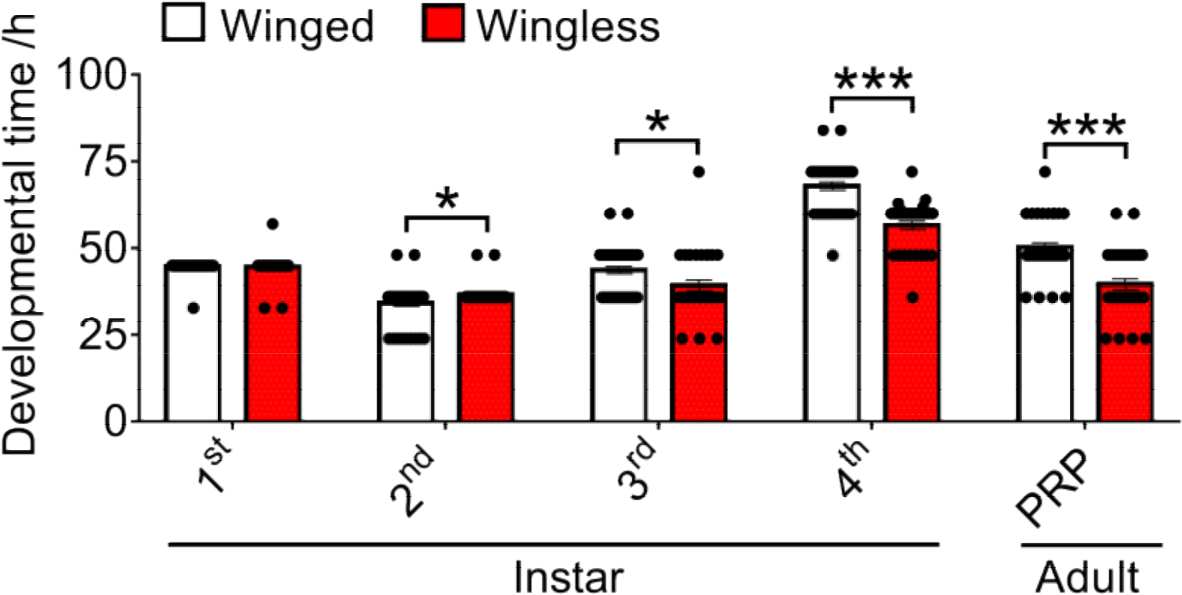
The developmental time of nymphal stages and pre-reproductive period (PRP) of winged- and wingless-destined aphids. Value in bar plots are means (±SE) of at least 36 replicates. Independent sample *t*-test was used to compare means, and significant differences between treatments are represented by asterisks: \**P*<0.05, \*\**P*<0.01, \*\*\**P*<0.001.

**Fig. S3.**
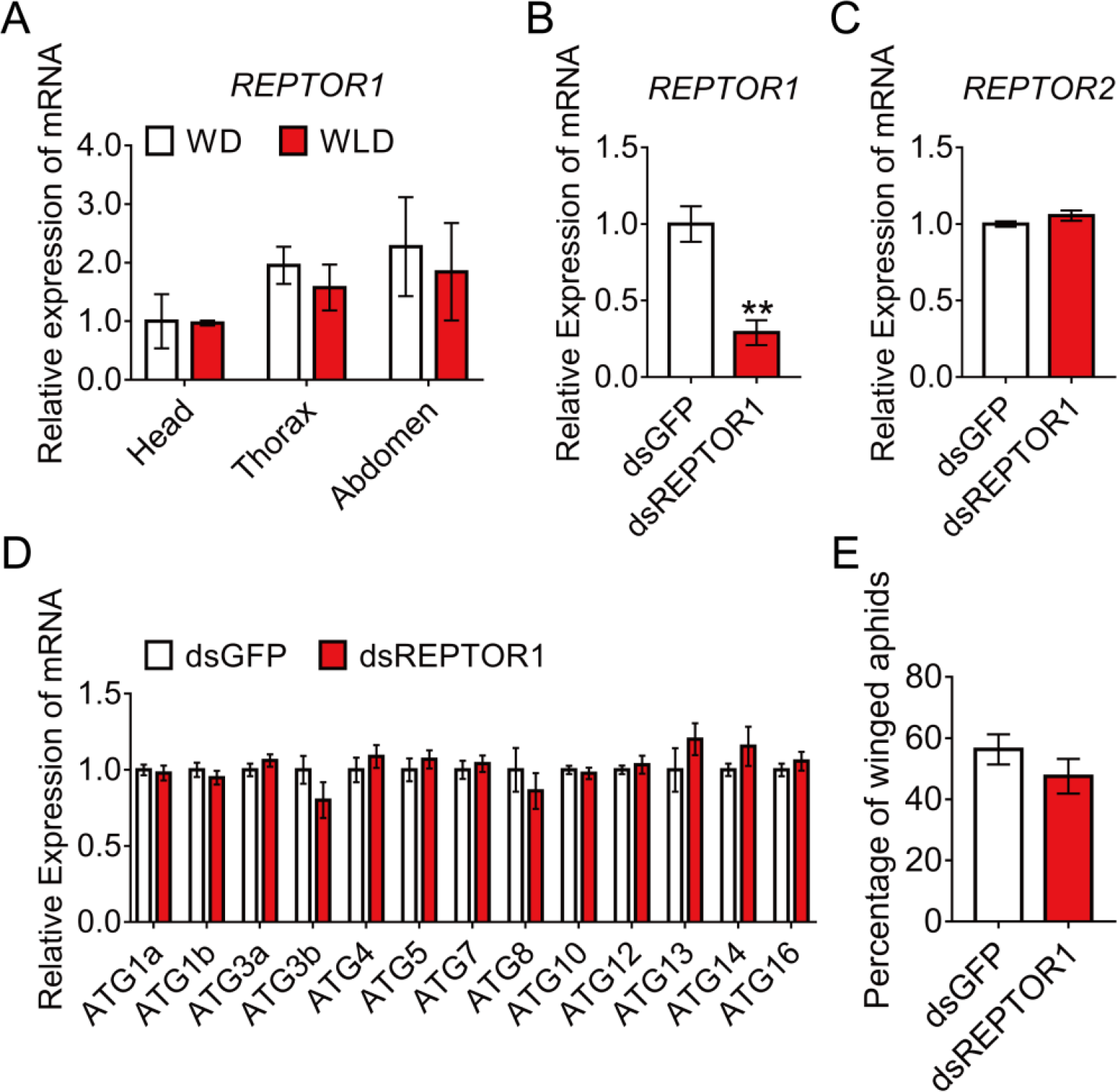
*REPTOR1* was not involved in activation of autophagic degradation in wing disc of winged-destined morphs. (A) Expression levels of *REPTOR1* in head, thorax and abdomen of winged- and wingless-destined aphids at 24 h post birth (n=4). (B-D) Knockdown of *REPTOR1* decreased the gene expression of *REPTOR1*, but did not affect the gene expression of *REPTOR2* and *ATG*s in whole body of the 1^st^ instar nymphs, as determined by qPCR (n=4). (E) *dsREPTOR1* did not affect the proportion of winged aphids (n=11). Independent sample t-test was used to compare means. Significant differences between treatments are represented by asterisks: \*\**P*<0.01.

**Fig. S4.**
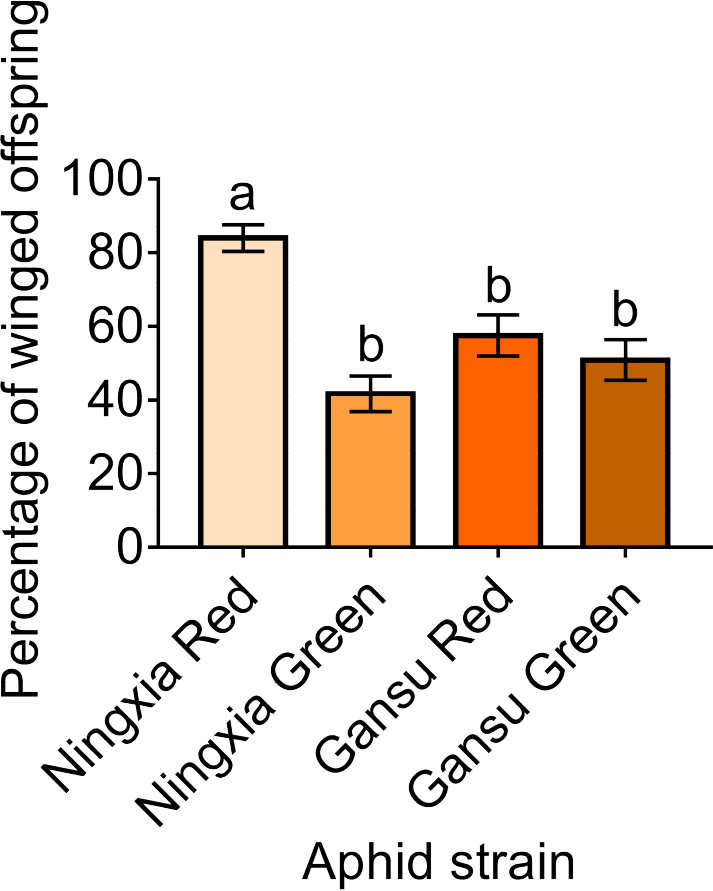
The percentage of winged offspring produced by two-adult contacting treatment for 4 h of different pea aphid strains. Values in bar represent means (±SE) of 20 replicates. Tukey’s multiple range tests at *P*<0.05 were used to compare means, and different lowercase letters indicate significant differences.

**Table S1.**
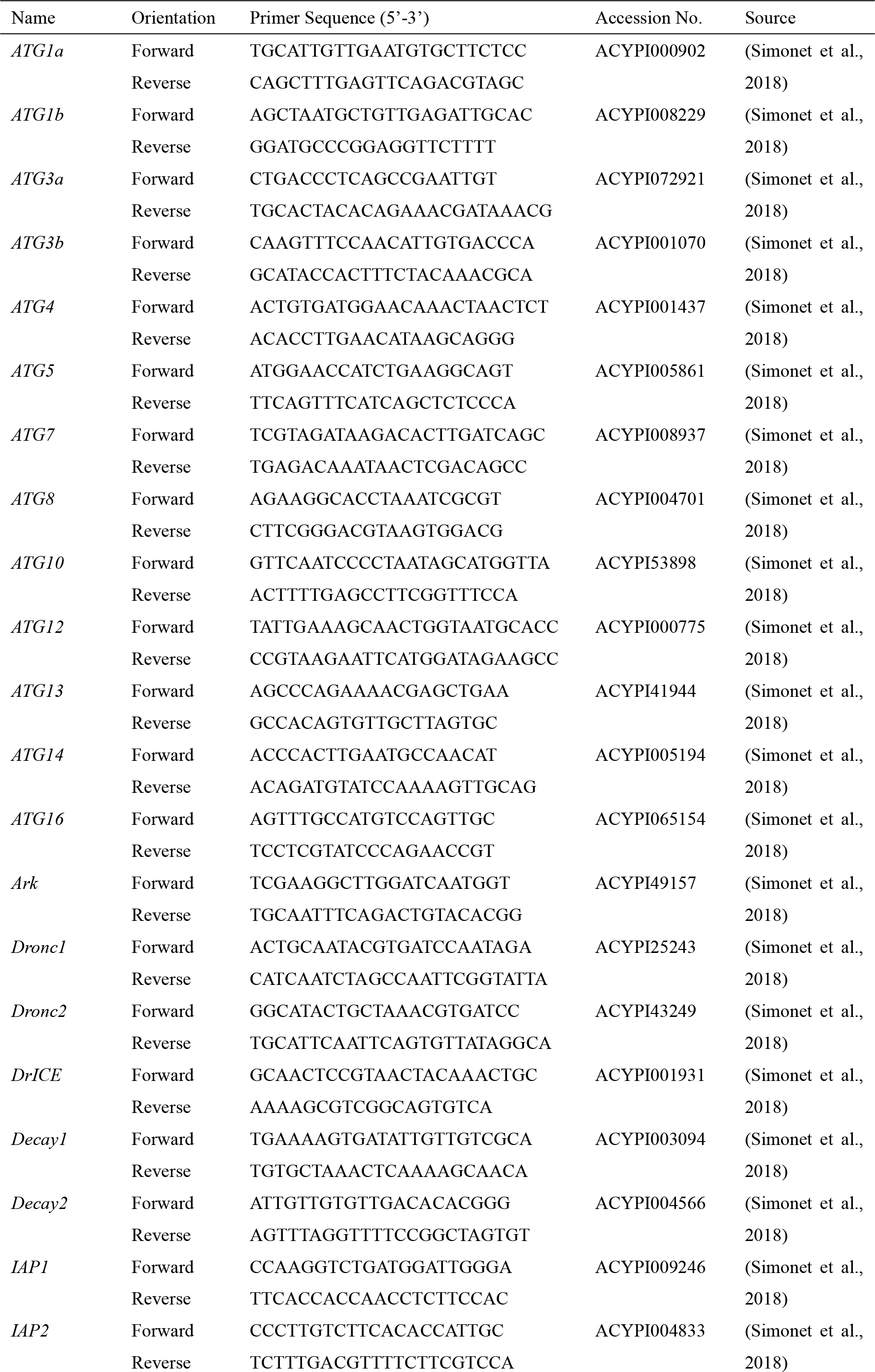
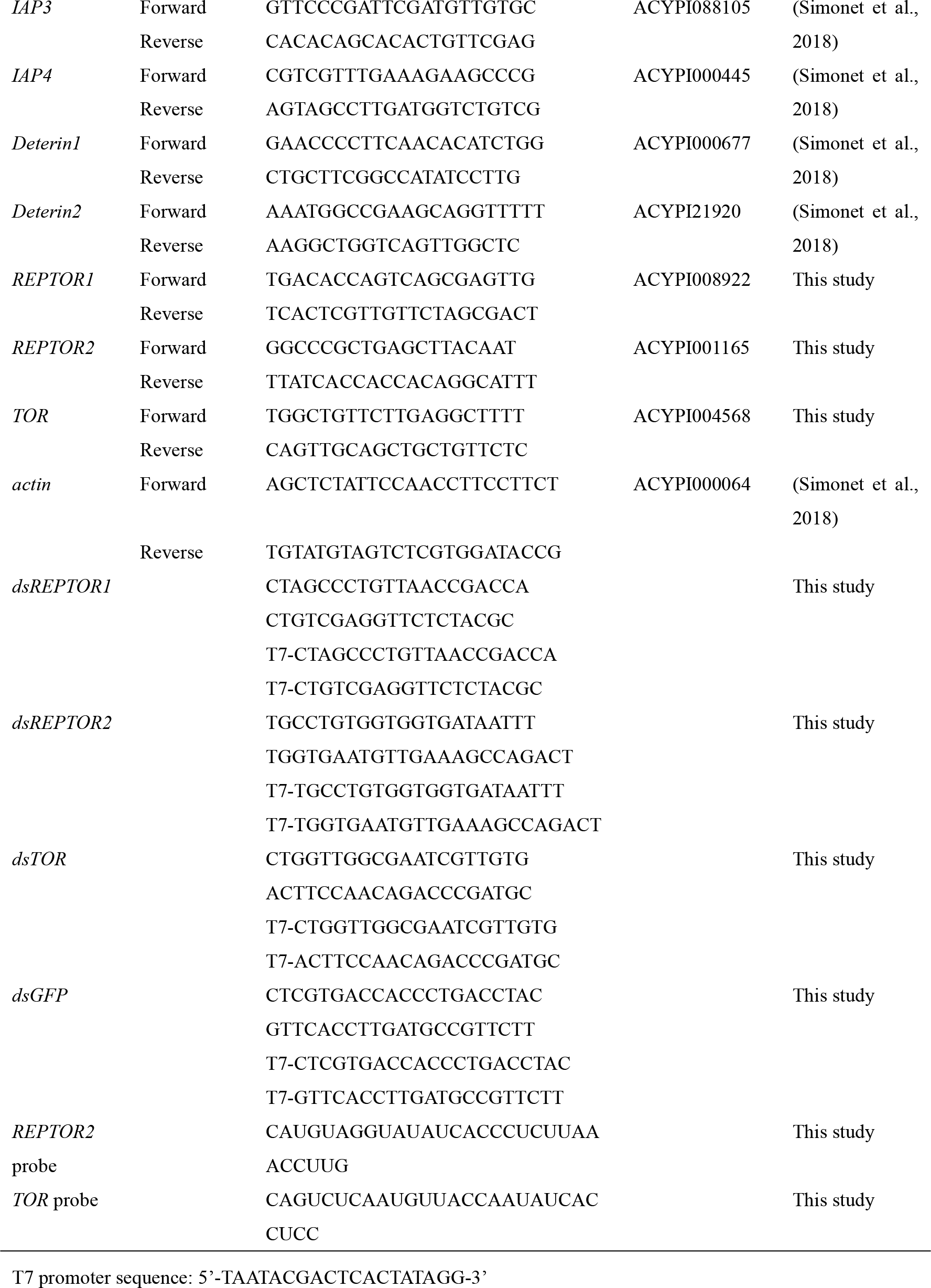
Primer List.

